# A chromosome-level genome assembly of the woolly apple aphid, *Eriosoma lanigerum* (Hausmann) (Hemiptera: Aphididae)

**DOI:** 10.1101/2020.05.29.121947

**Authors:** Roberto Biello, Archana Singh, Cindayniah J. Godfrey, Felicidad Fernández Fernández, Sam T. Mugford, Glen Powell, Saskia A. Hogenhout, Thomas C. Mathers

## Abstract

Woolly apple aphid (WAA, *Eriosoma lanigerum* Hausmann) (Hemiptera: Aphididae), is a major pest of apple trees (*Malus domestica*, order Rosales) and is critical to the economics of the apple industry in most parts of the world. Here, we generated a chromosome-level genome assembly of WAA – representing the first genome sequence from the aphid subfamily Eriosomatinae – using a combination of 10X Genomics linked-reads and *in vivo* Hi-C data. The final genome assembly is 327 Mb, with 91% of the assembled sequences anchored into six chromosomes. The contig and scaffold N50 values are 158 kb and 71 Mb, respectively, and we predicted a total of 28,186 protein-coding genes. The assembly is highly complete, including 97% of conserved arthropod single-copy orthologues based on BUSCO analysis. Phylogenomic analysis of WAA and nine previously published aphid genomes, spanning four aphid tribes and three subfamilies, reveals that the tribe Eriosomatini (represented by WAA) is recovered as a sister group to Aphidini + Macrosiphini (subfamily Aphidinae). We identified syntenic blocks of genes between our WAA assembly and the genomes of other aphid species and find that two WAA chromosomes (El5 and El6) map to the conserved Macrosiphini and Aphidini X chromosome. Our high-quality WAA genome assembly and annotation provides a valuable resource for research in a broad range of areas such as comparative and population genomics, insect-plant interactions and pest resistance management.

## 1 INTRODUCTION

There are approximately 5,000 known species of aphid (Hemiptera: Aphididae), and approximately 100 of these are of significant agricultural economic importance (Blackman & Eastop, 2017). While aphid genomics research has mostly focussed on the subfamily Aphidinae (International Aphid Genomics Consortium, 2010; Y. Li, Park, Smith, & Moran, 2019; Mathers, 2020; Mathers et al., 2017; Mathers, Mugford, et al., 2020; Mathers, Wouters, et al., 2020; Nicholson et al., 2015; Thorpe et al., 2018; Wenger et al., 2016), a large and widespread group including many important pests (Blackman & Eastop, 2000), investigation of genome evolution across the wider diversity of aphids has been limited (but see Julca et al., 2020). This report represents the first complete genome sequence for a species from the Eriosomatinae subfamily which includes three potentially polyphyletic tribes (Eriosomatini, Fordini and Pemphigini), all of which are distantly related to members of Aphidinae (X. Li, Jiang, & Qiao, 2014; Nováková et al., 2013; Ortiz-Rivas & Martínez-Torres, 2010).

Woolly apple aphid (WAA, *Eriosoma lanigerum* Hausmann, tribe Eriosomatini) is probably North American in origin. It was likely introduced to Britain in 1796 or 1797 with infested apple trees imported from America (Theobald, 1920) and has subsequently become a cosmopolitan and highly-damaging pest of apple worldwide (Barbagallo et al., 1997). Up to twenty generations per year on apple have been reported. First-instar nymphs (crawlers) are dispersive and walk to new feeding sites, but later developmental stages tend to be more sedentary, forming colonies distinguishable based on their fluffy white protective wax coating (Barbagallo et al., 1997). The aphid is able to feed on apple roots, trunks, branches and shoots. Saliva secreted whilst feeding causes cambium cells to divide rapidly, forming a gall which collapses and becomes pulpy under pressure from proliferating cells, making the area vulnerable to fungal infections (Barbagallo et al., 1997; Childs, 1929; Staniland, 1924). Edaphic (soil-dwelling) WAA has a significant negative effect on plant growth, especially of young, non-bearing apple trees where feeding significantly reduces trunk diameter (Brown, Schmitt, Ranger, & Hogmire, 1995) that is strongly correlated with fruit production (Waring, 1920). Such widespread and significant damage has made WAA resistance a key objective for rootstock breeding since the early 20th century (Cummins & Aldwinckle, 1983).

Sexual and asexual reproduction in aphid life cycles have essential impacts on population structure and allelic diversity (Delmotte, Leterme, Gauthier, Rispe, & Simon, 2002). Species in the subfamily Eriosomatinae typically go through phases of both sexual and parthenogenetic reproduction during the life cycle, but (unlike in other aphid subfamilies) the sexual males and egg-laying females lack mouthparts and are therefore unable to feed. The life cycle and mode of reproduction of WAA is somewhat ambiguous. In North America, where WAA populations have been reported to induce leaf galls on American elm (*Ulmus americana* L.) (Baker, 1915), it has been suggested that WAA has a heteroecious life cycle, alternating between apple (parthenogenetic reproduction) and elm (sexual reproduction). However, it is not clear whether genotypes found on elm are also capable of feeding on apple (Blackman & Eastop, 2000). In other parts of the world, WAA is assumed to be entirely anholocyclic (asexual) on apple (e.g. Dumbleton & Jeffreys, 1938; Eastop, 1966). Sexual males and females have sometimes been observed on apple, but the deposited eggs usually do not hatch, and such populations are assumed to be functionally asexual (Blackman & Eastop, 2000).

The genetic structure of WAA populations also has important practical relevance in the context of host-plant resistance. Four genes associated with WAA resistance in apple (*Er1-4*) have been identified from various sources (Bus et al., 2010; King et al., 1991; Ranatunga, Gardiner, Bssett, Rikkerink, & Bus, 1999). Some genotypes of WAA have been observed to feed on rootstocks with *Er1*-, *Er2*-, and *Er3*-mediated resistance (Cummins & Aldwinckle, 1983; Rock & Zeiger, 1974; Sandanayaka, Bus, Connolly, & Newcomb, 2003). The prevalence and spread of such resistance-breaking genotypes within WAA populations has not yet been investigated, and the availability of a full genome sequence will benefit such studies considerably.

Here, we generated a high-quality chromosome-level genome assembly of WAA using a combination of 10X Genomics linked-reads and *in vivo* chromatin conformation capture (Hi-C) sequencing. Subsequently, gene prediction, functional annotation and phylogenetic analysis were carried out to determine the relationship of WAA within the Aphidoidea superfamily. Our reference genome can provide information about genome organization in the Eriosomatinae subfamily and allows comparative genomic studies for a better understanding of the evolution of aphids.

## 2 MATERIALS AND METHODS

### 2.1 Sampling

Aphids were collected from a population feeding on apple trees grown under glass at NIAB EMR. The glasshouse-grown plants were deliberately infested using WAA collected from local orchards. A single colony infesting a glasshouse-grown potted apple plant (a clonally-propagated aphid-susceptible breeding line in the rootstock breeding programme at NIAB EMR) was sampled for aphids in November 2018. All sampled aphids were collected from one (12 cm) section of infested woody stem. Wax filaments covering sampled insects were removed using a fine paint brush and the aphids placed in Eppendorf tubes. Adult individuals (20 in total) were collected into separate tubes for genome analysis. An additional three samples of grouped aphids (25 in total) were pooled into individual tubes for RNA-Seq analysis. These groups consisted of: apterous (wingless) adults; mid-instar nymphs (mix of second, third and fourth instars); mixed stages (all of the above life stages present). Collected aphids were snap frozen by immersing tubes in liquid nitrogen.

### 2.2 DNA extraction and sequencing

The total genomic DNA was extracted from a single aphid using Illustra Nucleon Phytopure kit modifying the manufacturer’s protocol (GE Healthcare). The lysis step was performed adding 10 ul of Proteinase K and incubating the sample in a water bath for 2 hours at 55°C. The DNA precipitation was performed using NaAc (3M) together with Isopropanol for increasing the DNA yield. DNA quality and quantity were assessed using a Nanodrop spectrophotometer (Thermo Scientific), a Qubit double-stranded DNA BR Assay Kit (Invitrogen, Thermo Fisher Scientific) and Femto fragment analyser (Agilent). 10X Genomics library preparation and Illumina genome sequencing (HiSeq X, 150bp paired-end) were performed by Novogene Bioinformatics Technology Co, Beijing, China, in accordance with standard protocols.

A pool of mixed stages samples was used to construct a Hi-C chromatin contact map to enable a chromosome-level assembly. Dovetail Genomics created the Hi-C library with the DpnII restriction enzyme following a similar protocol to Lieberman-Aiden et al. (2009). Hi-C library was sequenced on Illumina HiSeq X sequencer and 150 bp paired-end reads were generated.

### 2.3 Genome assembly

To create the *de novo* assembly, the 10X Genomics linked-read data were assembled using Supernova 2.1.1 (Weisenfeld, Kumar, Shah, Church, & Jaffe, 2017) with the default parameters and the recommended number of reads (--maxreads=199222871) to produce the pseudohaplotype assembly output (--style=pseudohap). We improved the initial Supernova assembly by performing iterative scaffolding using all of the 10X Genomics raw data (364 millions of reads; **Supplementary Table 1**) following the procedure set out in Mathers, Wouters, et al. (2020). Briefly, we performed two rounds of Scaff10x (https://github.com/wtsi-hpag/Scaff10X) with the parameters “-longread 1 -edge 50000 - block 50000”, followed by mis-assembly detection and correction with Tigmint (Jackman et al., 2018). These steps were followed by a final round of scaffolding with ARCS (Yeo, Coombe, Warren, Chu, & Birol, 2018). We aligned the Hi-C reads to the 10x Genomics assembly using the Juicer pipeline (Durand et al., 2016). The assembly was then scaffolded with Hi-C data (**Supplementary Table 1**) into chromosome-level organization using the 3D-DNA pipeline (Dudchenko et al., 2017), followed by manual correction using Juicebox Assembly Tools (JBAT) (Dudchenko et al., 2018). The assembly was polished after JBAT review using the 3D-DNA seal module to reintegrate genomic content removed from super-scaffolds through false-positive manual editing to create a final scaffolded assembly.

We checked the Hi-C assembly for contamination using the BlobTools pipeline v0.9.19 (Kumar, Jones, Koutsovoulos, Clarke, & Blaxter, 2013; Laetsch & Blaxter, 2017) by generating taxon annotated GC content-coverage plots (known as “BlobPlots”). Each scaffold was annotated with taxonomy information based on BLASTN v2.2.31 (Camacho et al., 2009) searches against the NCBI nucleotide database (nt, downloaded 13/10/2017) with the options “-outfmt ‘6 qseqid staxids bitscore std sscinames sskingdoms stitle’ -culling_limit 5 -evalue 1e-25”. To calculate average coverage per scaffold, we mapped the 10X Genomics raw reads, after barcode removal using proc10xG (https://github.com/ucdavis-bioinformatics/proc10xG), to the assembly using BWA-MEM v0.7.7 (Li, 2013) with default parameters. The resulting BAM file was sorted with SAMtools v1.3 (Li et al., 2009) and passed to BlobTools along with the table of BLASTN results. The mitochondrial genome was identified and removed based on alignment to the WAA mitochondrial genome (NCBI accession number NC_033352.1) with nucmer v4.0.0.beta2 (Marçais et al., 2018), and patterns of coverage and GC content obtained from BlobTools. A frozen release was generated for the final assembly with scaffolds renamed and ordered by size with SeqKit v0.9.1 (Shen, Le, Li, & Hu, 2016). We assessed the quality of the genome assembly by searching for conserved, single copy, arthropod genes (n=1,066) with Benchmarking Universal Single-Copy Orthologs (BUSCO) v3.0 (Waterhouse et al., 2018) and by analysis of k-mer spectra with KAT v2.3.2 (Mapleson, Garcia Accinelli, Kettleborough, Wright, & Clavijo, 2017) using the default k-mer size (k = 27). To generate a k-mer spectra we compared k-mer content of the raw sequencing reads to the k-mer content of the assembly using KAT comp. Using this spectra we also estimated the WAA genome size, heterozygosity level and genome assembly completeness.

### 2.4 RNA extraction and sequencing

Total RNA was extracted from two pools of approximately 25 aphids, one containing adults and one containing mid-instar nymphs, collected from the same colony. The sample was ground under liquid nitrogen in a 1.5 ml Eppendorf tube using a plastic pestle. RNA was extracted using Trizol (Sigma) according to the manufacturer’s protocol. RNA was further purified using RNeasy with on-column DNAse treatment (Qiagen) according to the manufacturer’s protocol and eluted in 100 ml of nuclease-free water. RNA quality was assessed by electrophoresis of 5 μl denatured in formamide on a 1% agarose gel. Purity was assessed using a Nanodrop spectrophotometer (ThermoFisher) to measure the A_260_/A_280_ and A_260_/A_230_ ratios. Concentration of RNA was measured, and the presence of contaminating DNA was assessed using a Qubit (Lifetech).

Quality control and trimming for adapters and low-quality bases (quality score < 30) of the RNA-seq raw reads were performed using FastQC v0.11.8 (Andrews, 2010) and trim_galore v0.5.0 (http://www.bioinformatics.babraham.ac.uk/projects/trim_galore) respectively.

### 2.5 Gene prediction

Before running the gene prediction, we identified repeats and transposable elements (TEs) with RepeatMasker v4.0.7 (Tarailo-Graovac & Chen, 2009) using the RepBase Insecta repeat library (Bao, Kojima, & Kohany, 2015) with the parameters “-e ncbi -species insecta -a -xsmall -gff” (Jurka et al., 2005).

We mapped the quality and adapter trimmed RNA-seq reads (**Supplementary Table 2**) to the soft-masked assembly with HISAT2 v2.0.5 (Kim, Langmead, & Salzberg, 2015) with the following parameters: “--max-intronlen 25000 --dta-cufflinks” followed by sorting and indexing with SAMtools v1.3 (Li et al., 2009). Strand-specific RNA-seq alignments were split by forward and reverse strands and passed to BRAKER2 as separate BAM files. Therefore, we ran BRAKER2 v2.1.2 (Hoff, Lange, Lomsadze, Borodovsky, & Stanke, 2016; Hoff, Lomsadze, Borodovsky, & Stanke, 2019) to train AUGUSTUS (Lomsadze, Burns, & Borodovsky, 2014; Stanke, Diekhans, Baertsch, & Haussler, 2008) and predict protein-coding genes, incorporating evidence from the RNA-seq alignments and alignment of BUSCO genes with the following parameters “--softmasking --gff3 --prg=gth --gth2traingenes”. After gene prediction, completeness of the gene set was checked with BUSCO using the longest transcript of each gene as the representative transcript.

### 2.6 Functional annotation

All the unique transcripts were converted to peptide sequence using cufflinks v2.2.1 (Trapnell et al., 2010). Sequences were searched against the non-redundant NCBI protein database using BLASTP v2.6.0 with an E-value cut-off of <= 1×10^−5^. BLAST2GO v5.0 (Conesa et al., 2005) and InterProScan v2.5.0 (Quevillon et al., 2005) were used to assign Gene Ontology (GO) terms. Protein domains were annotated by searching against the InterPro v32.0 (Hunter et al., 2012) and Pfam v27.0 (Punta et al., 2012) databases, using InterProScan v2.5.0 (Quevillon et al., 2005) and HMMER v3.1 (Finn, Clements, & Eddy, 2011), respectively. The pathways in which the genes might be involved were assigned by BLAST against the KEGG databases (release 53), with an E-value cut-off of 1×10^−5^.

### 2.7 Phylogeny and comparative genomics

Orthologous groups in Aphididae genomes were identified from the predicted protein sequences of WAA and nine other aphid genomes already published (**Supplementary Table 3**): *Myzus cerasi* (Mathers, Mugford, et al., 2020; Thorpe et al., 2018), *Myzus persicae* (Mathers, Wouters, et al., 2020), *Diuraphis noxia* (Nicholson et al., 2015), *Acyrthosiphon pisum* (Mathers, Wouters, et al., 2020), *Pentalonia nigronervosa* (Mathers, Mugford, et al., 2020), *Aphis glycines* (Mathers, 2020), *Rhopalosiphum maidis* (Chen et al., 2019), *Rhopalosiphum padi* (Thorpe et al., 2018) and *Cinara cedri* (Julca et al., 2020). As an outgroup, we included the genome of the silverleaf whitefly *Bemisia tabaci* (Chen et al., 2016). We used the longest transcript to represent the gene model when several transcripts of a gene were annotated. OrthoFinder v2.2.3 (Emms & Kelly, 2015, 2019) with Diamond v0.9.14 (Buchfink, Xie, & Huson, 2015), MAFFT v7.305 (Katoh & Standley, 2013) and FastTree v2.1.7 (Price, Dehal, & Arkin, 2009, 2010) were used to cluster proteins into orthogroups, reconstruct gene trees and estimate the species tree. The OrthoFinder species tree was automatically rooted by OrthoFinder based on informative gene duplications with STRIDE (Emms & Kelly, 2017).

Gene Ontology (GO) analysis of lineage-specific gene families and genes that have undergone lineage-specific duplication was carried out using the Bioconductor (Gentleman et al., 2004) package topGO (Alexa & Rahnenführer, 2009). We used a Fisher exact test to identify overrepresented GO terms.

### 2.8 Synteny analysis

Syntenic blocks of genes were identified between the chromosome-level genome assemblies of WAA, *M. persicae, A. pisum* and *R. maidis* (see **Supplementary Table 3** for details of assembly and annotation versions used) using MCscanX v1.1 (Wang et al., 2012). For each comparison, we carried out an all vs. all BLAST search of annotated protein sequences using BLASTALL v2.2.22 (Altschul, Gish, Miller, Myers, & Lipman, 1990) with the options “-p blastp - e 1e-10 -b 5 -v 5 -m 8” and ran MCScanX with the parameters “-s 5 -b 2”, requiring synteny blocks to contain at least five consecutive genes and to have a gap of no more than 25 genes. MCScanX results were visualised with SynVisio (https://synvisio.github.io/#/).

## 3 RESULTS AND DISCUSSION

### 3.1 Genome sequencing and assembly

We generated a high-quality chromosome-level genome assembly of WAA using a combination of 10X Genomics linked-reads and *in vivo* Hi-C data (**Figure 1a**). In total, we generated 54.72 Gb of 10X Genomics linked-reads and 71.95 Gb of Hi-C reads, corresponding to 167x and 220x coverage of the WAA genome, respectively. Initial *de novo* assembly of the 10X Genomics linked-reads with Supernova produced a contiguous assembly totalling 330 Mb (**Table 1**; scaffold N50 = 4.16 Mb). We further improved the Supernova assembly by iterative scaffolding and misjoin correction with Scaff10X (two rounds) and Tigmint (one round), increasing the scaffold N50 of the assembly to 4.22 Mb and reducing the number of scaffolds from 8,967 to 7,929, with the longest scaffold spanning 12.58 Mb (**Table 1**). To generate chromosome length super-scaffolds, we scaffolded the 10X assembly using *in vivo* Hi-C data. After manual curation, the final assembly comprised 327 Mb, with 91% of the assembly anchored into six chromosomes (**Figure 1a**), consistent with the WAA 2n karyotype being 12 chromosomes (Gautam & Verma, 1982, 1983; Kulkarni, 1984; Robinson & Chen, 1969). The lengths of the six chromosomes ranged from 29.68 to 71.23 Mb.

**Figure 1.**
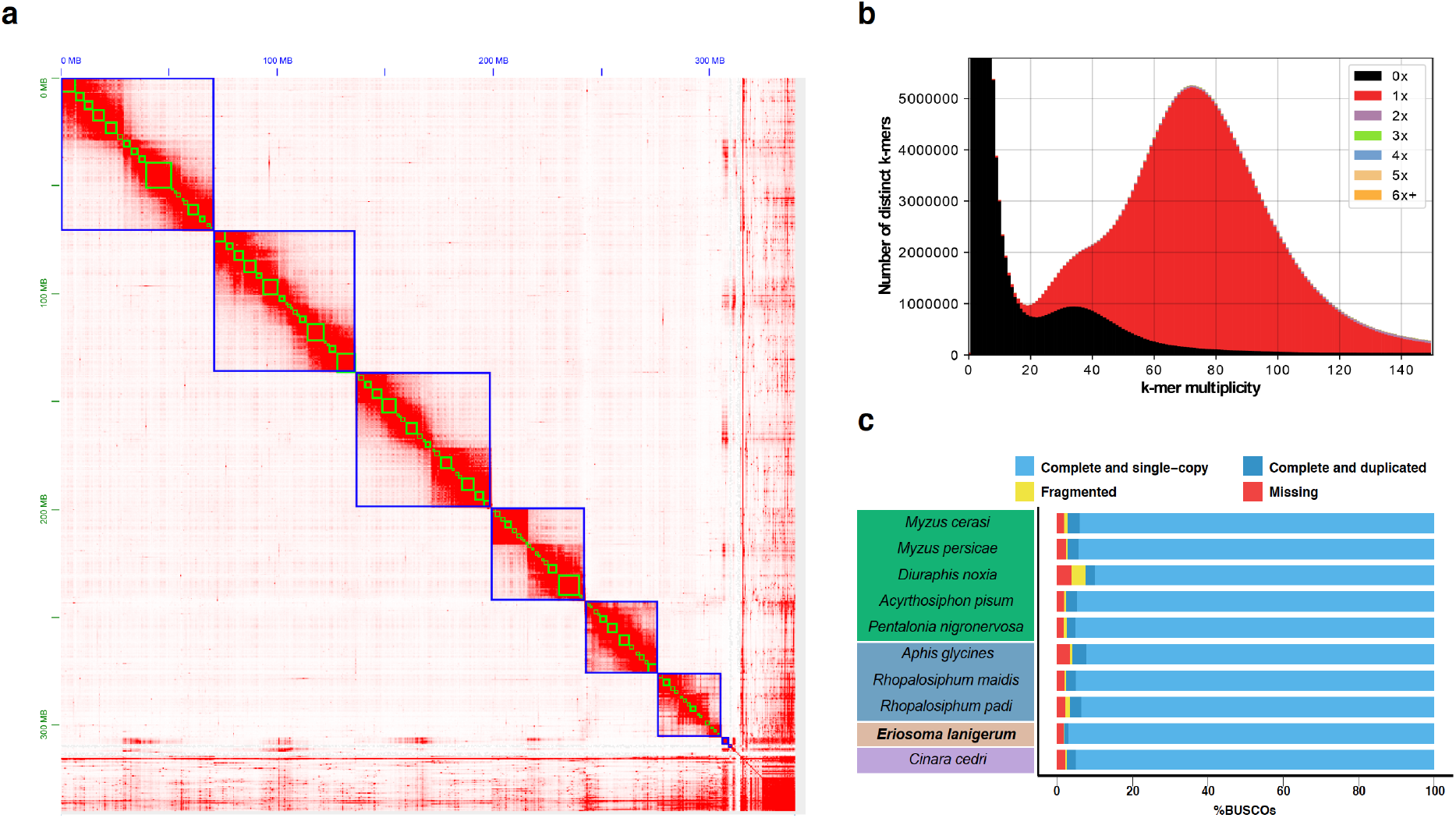
Chromosome-level genome assembly of WAA. (**a**) Heatmap showing frequency of Hi-C contacts along WAA genome assembly. Blue lines indicate super scaffolds and green lines show contigs. Genome scaffolds are ordered from longest to shortest with the X and Y axis showing cumulative length in millions of base pairs (Mb). (**b**) KAT k-mer plots comparing k-mer content of 10X Genomics raw reads (barcodes removed) with Hi-C assembly. The black area of the graphs represents the distribution of k-mers present in the reads but not in the assembly and the red area represents the distribution of k-mers present in the reads and once in the assembly. Other colours show k-mers found multiple times in the genome assembly. (**c**) Genome assembly completeness assessed by the recovery of universal single-copy genes (BUSCOs) using the Arthropoda gene set (n=1,066). BUSCO assessment result for *Eriosoma lanigerum* (in bold) compared with aphid genomes available from National Center for Biotechnology Information (NCBI). The species are coloured by aphid tribe (see **Figure 2**).

**Figure 2.**
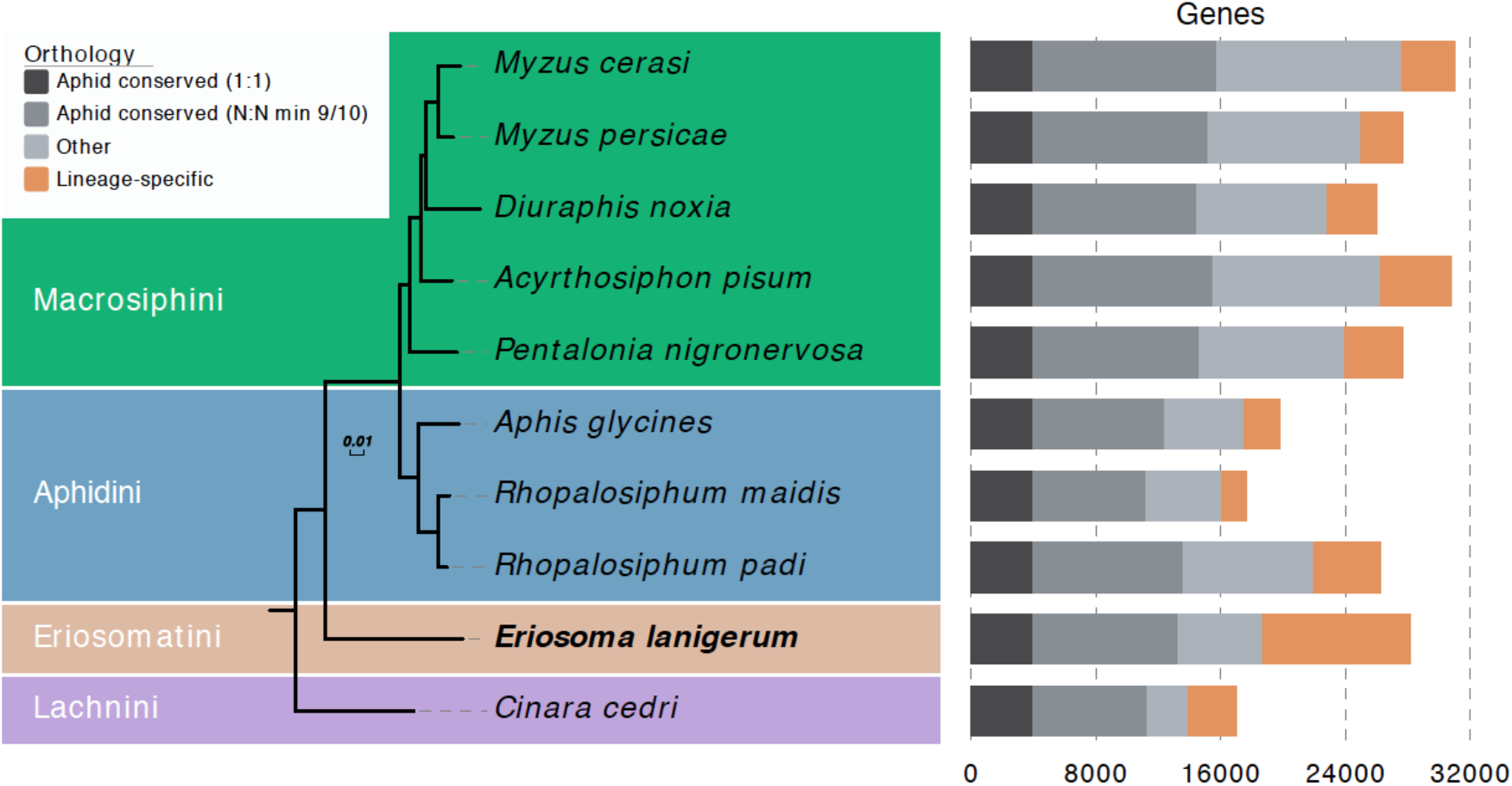
Maximum likelihood phylogeny of *Eriosioma lanigerum* and other nine aphid species based on a concatenated alignment of 3,079 conserved one-to-one orthologs. The tree is rooted with the whitefly *B. tabaci* MEAM1 (not shown). Clades are coloured by aphid tribe. Branch lengths are in amino acid substitutions per site. The *bar chart* shows the gene count for each species with orthology relationships among aphids.

**Table 1.** Genome summary statistics for each step of the assembly of WWA. SP = Supernova; SC = Scaff10X; TG = Tigmint; HC = Hi-C.

| *Assembly* | *SP* | *SP+SC+TG* | *SP+SC+TG+HC* |
| --- | --- | --- | --- |
| **Base pair (Mb)** | 330 | 330 | 327 |
| **Number of contigs** | 12,566 | 12,703 | 12,065 |
| **Contig N50 (Mb)** | 0.158 | 0.158 | 0.158 |
| **Number of scaffolds** | 8,967 | 7,929 | 7,146 |
| **Scaffold N50 (Mb)** | 4.164 | 4.222 | 62.861 |
| **Longest scaffold (Mb)** | 16.081 | 12.584 | 71.231 |
| **% of assembly in chromosome length scaffolds** | 0 | 0 | 91 |

The WAA genome assembly is accurate, complete and free from contamination. K-mer analysis comparing genomic content of the 10X reads (after barcode removal) with the WAA genome assembly reveals little missing single-copy genome content and very low levels of duplicated content caused by the assembly of haplotigs (**Figure 1b**). Furthermore, our 327 Mb WAA genome assembly is close to the genome size estimate based on k-mer analysis with KAT (363 Mb) (**Figure 1b**; **Supplementary Table 4**) and the genome size of *Eriosoma americanum* (330 Mb) (Finston, Hebert, & Foottit, 1995). This analysis is further supported by high representation of conserved arthropod genes (n=1,066) in the assembly, with 97% (n=1,032) found as complete single copies. Indeed, the WAA assembly contains the highest number of conserved single-copy Arthropoda genes of any published aphid genome (**Figure 1c**). A taxon-annotated GC content-coverage plot (known as a “BlobPlot”; Kumar, Jones, Koutsovoulos, Clarke, & Blaxter, 2013) revealed the co-assembly of the obligate aphid bacterial symbiont *Buchnera aphidicola* (Baumann et al., 1995; Douglas, 1998; Hansen & Moran, 2011; Shigenobu & Wilson, 2011) and a secondary symbiont, *Serratia symbiotica* (Burke & Moran, 2011; Manzano-Marín & Latorre, 2016; Moran, Russell, Koga, & Fukatsu, 2005) (**Supplementary Figure 1)**. These bacterial scaffolds were filtered from the final assembly, along with scaffolds showing atypical GC content and read coverage, leaving the final assembly free from obvious contamination (**Supplementary Figure 2)**. *B. aphidicola* genome was assembled into 99 scaffolds 444 Kb in length (**Supplementary Table 5**). *S. symbiotica* genome was fragmented and incomplete (165 Kb total length, 30 scaffolds, N50 = 9 Kb).

### 3.2 Genome annotation

We generated 44 Gb of strand-specific RNA-seq data to aid genome annotation. A total of 28,186 protein-coding genes (28,297 transcripts) were predicted in the WAA genome assembly. Completeness of the gene set reflected that of the genome assembly, with 97.3% of the BUSCO Arthropoda gene set found as complete copies (95.6% complete and single copy; **Supplementary Figure 3**). In total, 55.7% (15,477) of the predicted transcripts were functionally annotated with at least one GO-term and/or protein domain. 9,903 transcripts were assigned GO term annotations using BLAST2GO and 9,934 using InterProScan. 14,228 transcripts contained at least one known InterProScan protein domain.

### 3.3 Phylogeny and comparative genomics

WAA is the first member of the aphid subfamily Eriosomatinae to have its genome sequenced. To place this new genome assembly in a phylogenetic context and to investigate gene family evolution across aphids, we compared the WAA proteome (the complete set of annotated protein coding genes) to the proteomes of nine other aphid species that have fully sequenced genomes (**Supplementary Table 3**) and to the whitefly, *Bemisia tabaci* MEAM1 (Chen et al., 2016). In total, we clustered 240,702 proteins into 23,294 orthogroups (gene families) and 26,969 singleton genes (**Supplementary Table 6**). Maximum likelihood phylogenetic analysis based on a concatenated alignment of 3,079 conserved single-copy genes produced a fully resolved species tree with 100% support at all nodes (**Figure 2**). Eriosomatini (represented by WAA) is recovered as a sister group to Aphidini + Macrosiphini (Aphidinae), with Lachnini (represented by *C. cedri*) placed as an outgroup to all other sequenced aphid species (**Figure 2**).

The number of predicted genes in WAA (28,186) is within the range of other aphid genomes (16,992 – 31,001) but among the highest (**Figure 2**). However, predicted gene numbers can vary depending on both the quality of the genome assembly and the different pipelines used for predicting genes (Denton et al., 2014; Yandell & Ence, 2012). Of the 28,186 predicted genes in the WAA genome, 18,738 (66%) have an ortholog in at least one other aphid species and 13,278 (47%) are conserved in the majority (at least 9/10) of aphid species (**Figure 2**). The high number of genes with orthologs in other aphid species despite the absence of close WAA relatives in our analysis likely reflects the early emergence of many aphid gene families (Julca et al., 2020).

Aphid genomes are also subject to high levels of ongoing gene duplication (Fernández et al., 2020; International Aphid Genomics Consortium, 2010; Julca et al., 2020; Mathers et al., 2017; Thorpe et al., 2018). WAA is no exception and we detect a large number of lineage-specific gene families (689 orthogroups corresponding to 3,954 genes; **Figure 2**) and identify 9,936 genes that have undergone lineage-specific duplication. These genes are enriched for a diverse set of functions including sensory perception and metabolic process (**Supplementary Table 7**; **Supplementary Table 8**). As additional Eriosomatini genomes become available, the diversification of these genes and gene families will be investigated in greater detail.

### 3.4 X chromosome fragmentation in WAA

We have previously shown that the autosomes of aphids within the tribes Macrosiphini and Aphidini have undergone extensive rearrangement over the last ~30 million years while the aphid sex (X) chromosome has been conserved (Mathers, Wouters, et al., 2020). Given that we now have a chromosome-level genome assembly of an aphid from a third, more divergent tribe, we used our WAA aphid assembly to investigate the evolution of aphid genome structure. We identified syntenic blocks of genes between our WAA assembly and the genomes of *M. persicae* (**Figure 3a**) and *A. pisum* (**Figure 3b**) from Macrosiphini (Mathers, Wouters, et al., 2020), and *R. maidis* (**Figure 3c**) from Aphidini (Chen et al., 2019). All comparisons reveal high levels of genome rearrangement between the autosomes. Surprisingly, however, we find that two WAA chromosomes (El5 and El6) map to the conserved Macrosiphini and Aphidini X chromosome, suggesting either fragmentation of the X chromosome in WAA or that the large Aphidinae (Macrosiphini + Aphidini) X chromosome was the result of an ancient fusion event. Additional chromosome-level assemblies of diverse aphid species will be required to test these two competing hypotheses. However, the lack of autosomal rearrangements between either El5 or El6 and the autosomes, suggests that recent X chromosome fragmentation in WAA is more likely.

**Figure 3:**
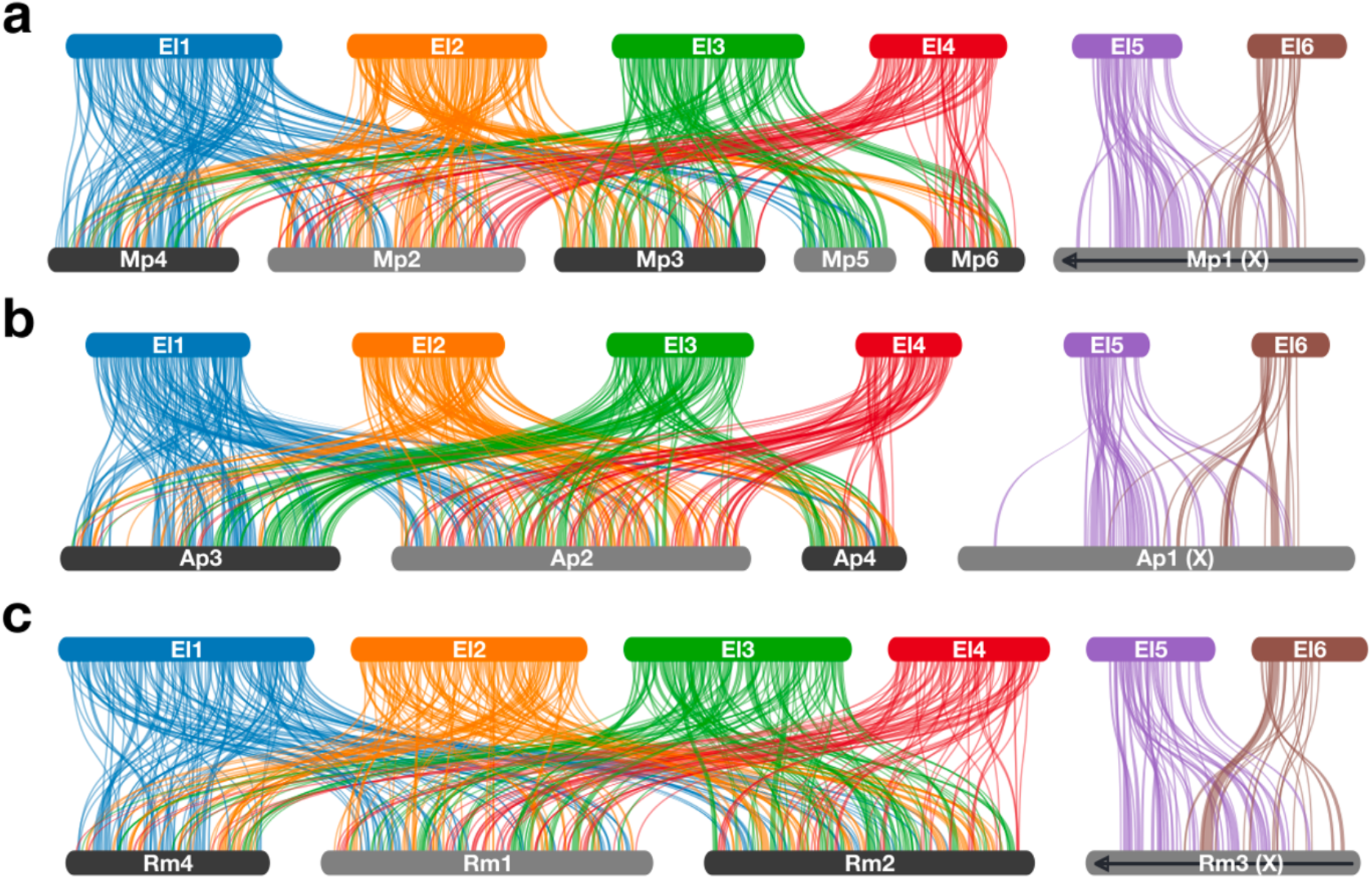
Genome reorganisation across aphids and fragmentation of the sex (X) chromosome in WAA. Pairwise synteny relationships are shown between WAA and the chromosome-scale genome assemblies of *Myzus persicae* (**a**), *Acyrthosiphon pisum* (**b**) and (**c**) *Rhopalosiphum maidis* (see **Figure 2** for phylogenetic relationships between compared species). Links indicate the boundaries of syntenic blocks of genes identified by MCscanX and are colour coded by WAA chromosome ID. Black arrows along chromosomes indicate reverse compliment orientation relative to WAA.

## 4 CONCLUSIONS

WAA is a widespread pest of apple trees that is particularly critical to the economics of the apple industry in most parts of the world. This study provides the first chromosome-level genome of WAA and the genome sequencing of the first representative of the whole subfamily Eriosomatinae. The WAA genome will be useful as a reference for investigation of genetic differences among wild WAA populations. The high quality of the genome will also allow molecular marker development and detection using large-scale genome resequencing. Population resequencing data can be used to investigate genomic regions and functional genes that show genetic variability and to analyse demographic history events in WAA populations. Finally, as the only genome available for the Eriosamatinae subfamily and as an additional outgroup to other sequenced aphids from the subfamily Aphidinae, the WAA genome will allow more extensive comparative genomics analysis of aphids.

## DATA ACCESSIBILITY

The genome raw reads, the RNA sequencing data and the genome assembly are available at the National Center for Biotechnology Information (NCBI) with the BioProject number PRJNA623270. The genome assembly and annotation and orthogroup clustering results are available for download from Zenodo (10.5281/zenodo.3797131).

## ACKNOWLEDGMENTS

TCM is funded by a BBSRC Future Leader Fellowship (BB/R01227X/1). Additional support was received from the BBSRC Institute Strategy Programme (BB/P012574/1) and the John Innes Foundation. CJG acknowledges receipt of a studentship funded by the BBSRC, AHDB and an industry consortium (http://www.ctp-fcr.org/industry-partners/).

## AUTHOR CONTRIBUTIONS

GP, SAH and TCM conceived the study. GP, FFF and CJG provided samples. RB and STM extracted DNA and RNA. RB, AS and TCM performed the genome assembly, gene model prediction, gene annotation, and comparative analyses. RB, CJG, GP, SAH and TCM wrote the manuscript with input from all authors. All authors reviewed the manuscript.

