## Supplementary Figure for "A chromosome-level genome assembly of the woolly apple aphid, *Eriosoma lanigerum* (Hausmann) (Hemiptera: Aphididae)"

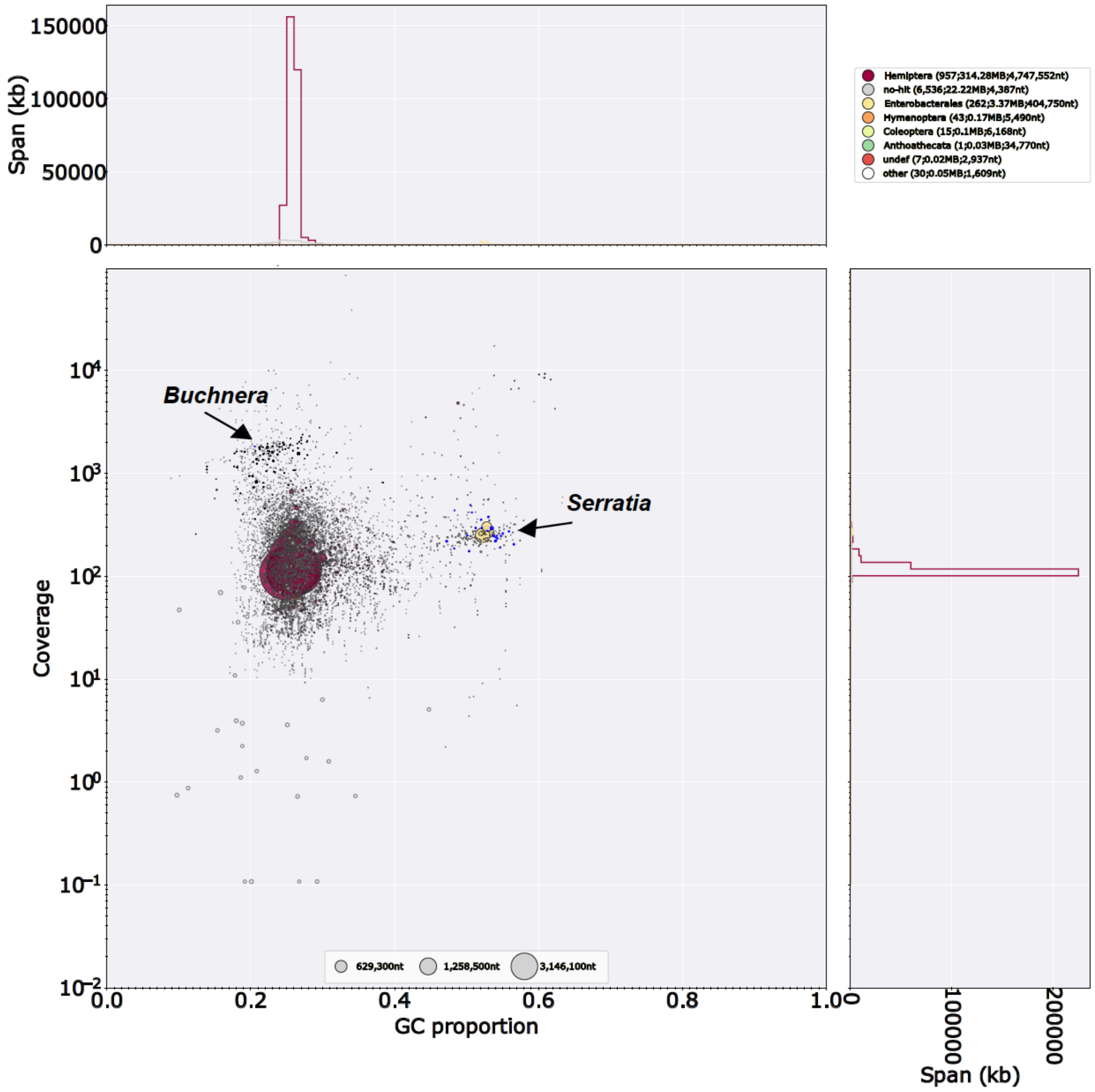


**Supplementary Figure 1**. Taxon-annotated GC content-coverage plot of WAA genome assembly before contaminants removing. Each circle represents a scaffold in the assembly, scaled by length, and coloured by order-level NCBI taxonomy assigned by BlobTools. The X axis corresponds to the average GC content of each scaffold and the Y axis corresponds to the average coverage based on alignment of raw 10X Genomics reads. Marginal histograms show cumulative genome content (in Kb) for bins of coverage (Y axis) and GC content (X axis). Arrows highlight scaffolds assigned to the symbiotic bacteria *Buchnera* (black circles) and *Serratia* (blue circles) which were removed from the final assembly.

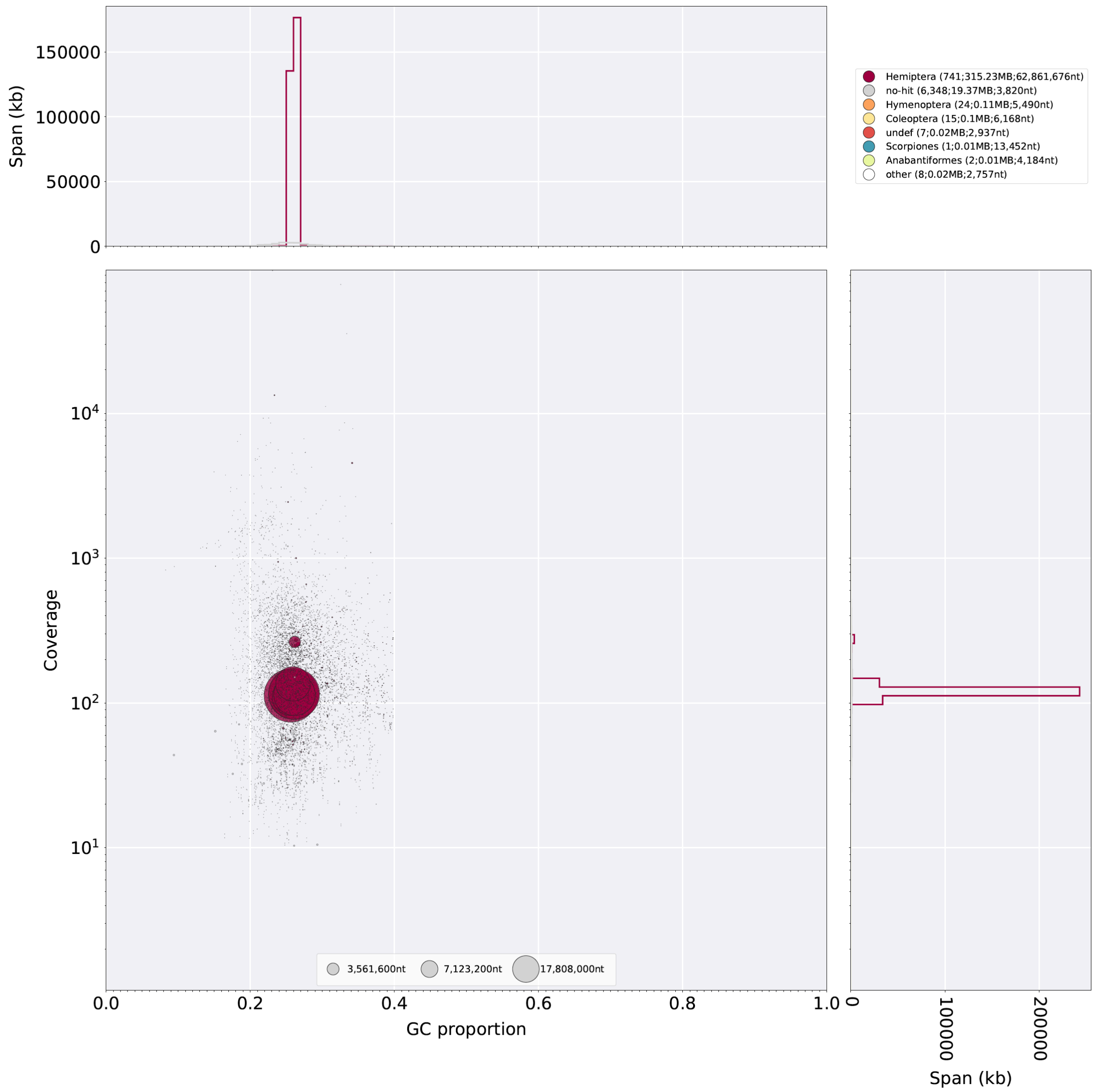


**Supplementary Figure 2**. Taxon-annotated GC content-coverage plot of WAA genome assembly after contaminants removing. Each circle represents a scaffold in the assembly, scaled by length, and coloured by order-level NCBI taxonomy assigned by BlobTools. The X axis corresponds to the average GC content of each scaffold and the Y axis corresponds to the average coverage based on alignment of raw 10X Genomics reads. Marginal histograms show cumulative genome content (in Kb) for bins of coverage (Y axis) and GC content (X axis).

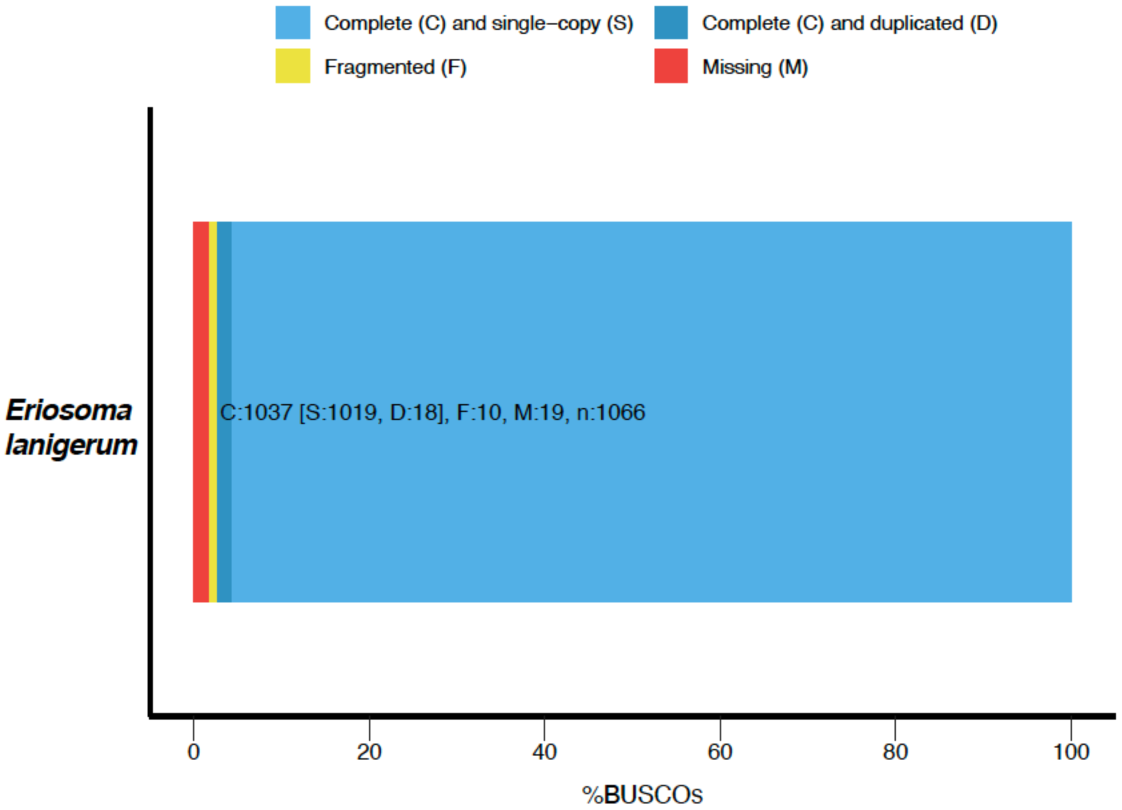


**Supplementary Figure 3**. BUSCO analysis of the *Eriosoma lanigerum* gene sets (protein sequences). Where multiple transcripts of a gene where annotated we used the longest transcript to represent the gene. The protein sets were assessed using the Arthropoda gene set (n=1,066).
